# Cell Segmentation in Images Without Structural Fluorescent Labels

**DOI:** 10.1101/2023.01.12.523757

**Authors:** Daniel Zyss, Susana A. Ribeiro, Mary J.C. Ludlam, Thomas Walter, Amin Fehri

## Abstract

High-content screening (HCS) provides an excellent tool to understand the mechanism of action of drugs on disease-relevant model systems. Careful selection of fluorescent labels (FLs) is crucial for successful HCS assay development. HCS assays typically comprise (1) FLs containing biological information of interest, and (2) additional structural FLs enabling instance segmentation for downstream analysis. However, the limited number of available fluorescence microscopy imaging channels restricts the degree to which these FLs can be experimentally multiplexed. In this paper, we present a segmentation workflow that overcomes the dependency on structural FLs for image segmentation, typically freeing 2 fluorescence microscopy channels for biologically relevant FLs. It consists in extracting structural information encoded within readouts that are primarily biological, by fine-tuning pre-trained state-of-the-art generalist cell segmentation models for different combinations of individual FLs, and aggregating the respective segmentation results together. Using annotated datasets that we provide, we confirm our methodology offers improvements in performance and robustness across several segmentation aggregation strategies and image acquisition methods, over different cell lines and various FLs. It thus enables the biological information content of HCS assays to be maximized without compromising the robustness and accuracy of computational single-cell profiling.

**Impact Statement:** This methodological paper describes a framework enabling cell segmentation for datasets without structural fluorescent labels to highlight cell organelles. Such capabilities favorably impact costs and possible discoveries in single-cell downstream analysis by improving our ability to incorporate more biological read-outs into a single assay. The perspective of computational and experimental biologist coauthors ensures a multidisciplinary viewpoint and accessibility for a wide readership.

## 1. Introduction

Image-based cellular assays allow us to investigate cellular and population phenotypes and signaling and thus to understand biological phenomena with high precision. In order to investigate specific biological processes and pathways, one can use tailored *fluorescent labels* (FL) such as immunofluorescence staining or fluorescent proteins. Cellular and subcellular features can then be detected and quantified by measuring fluorescence signal intensity and localization or by using multi-parametric measurements in a machine-learning framework [1]. Cell instance segmentation is a key part of such bioimage analysis pipelines as it allows us to study the cells at a single-cell level rather than at the population level.

Each FL has a function in an assay design. If a FL has as a primary function to label a specific cellular compartment, we call it a *structural* FL; otherwise it is a *non-structural* FL. Some non-structural FLs tend to consistently label cellular compartments, even if it is not their primary function. We refer to them as *structurally strong*. Indeed, a FL can also, on top of its primary role (e.g. to label a protein associated with a signaling pathway), highlight cellular structures useful for segmentation. If they do not consistently do so, we call them *structurally weak*. For example, due to the unique behavior of individual cells when exposed to chemical compounds, signaling pathways highlighted by non-structural FLs can lead to translocation to different cellular compartments, increased/decreased expression, or altered distribution. In fact, we observe that non-structural FLs lie on a continuum between both categories: the structural and morphological information they carry can vary significantly.

Deep learning models for cell instance segmentation have recently reached the quality of manual annotations [2]–[4], especially thanks to the emergence of models like U-Net [5]. Cellpose [4] is a notable U-Net based approach, which uses a multi-modal training dataset spanning several cell types and cell lines imaged under a variety of different imaging methods. It also benefits from being associated with a large community which incrementally increases the size and diversity of the dataset, in turn improving the performance of the model. Cellpose approaches the problem of multiple instance segmentation by predicting spatial gradient maps from the images, from which individual cell segmentations can be inferred. Other recent deep learning segmentation methods for cell biology are StarDist [2] and NucleAIzer [3] which make use of the U-Net architecture as well but with different representation for their images, and Mask-RCNN [6] which directly segments regions of interests (ROIs) in images with deep learning methods. We select Cellpose as a foundation for our approach as it is a widely used and well-designed framework which offers to be the most generalist with its community driven, ever-expanding training dataset.

However, a limitation of Cellpose is its heavy reliance on structural FLs for nucleic and cytoplasmic segmentation, predominantly FLs such as DAPI [7] for nuclei and Nuclear Export Signal (NES) [8] for cytoplasm. Indeed, only 15% of the Cellpose training dataset contain other fluorescent FLs. While such structural FLs are generally integrated in assay designs, being able to segment cells without using them allows to maximise the number of non-structural FLs. Since the total number of available microscopy channels is subject to numerous limitations – such as the bleed through effect [9] – and is typically limited to 4, the two extra channels made available can lead to assays delivering richer information about cellular processes.

In this paper, we demonstrate that non-structural FLs may contain information about cell and nuclear morphology that can be leveraged, even though they are not optimized in this regard, in contrast with structural FLs. Additionally we demonstrate that a collection of non-structural FLs are a sufficient substitute for structural FLs with regards to segmentation.

We note that our approach follows recent trends towards the “expertization” of generalist models like Cellpose 2.0 [10], encouraging the prediction of a wider range of cellular image types and styles, with very small additional training from humans in the loop. However, while those recent approaches still only apply to cell images with structural FLs, we hereby propose an extension to non-structural FLs.

We thus propose a generic framework for nuclear and cytoplasmic segmentation without the need to include corresponding structural FLs in the assay design. Our framework requires few annotations to finetune a pre-trained generalist deep learning segmentation base-model –here Cellpose– on each FL with a small set of annotated images. We show, on multiple datasets that we provide, that by combining the predictions from multiple non-structural FLs with various structural characteristics, we are able to reach segmentation performance comparable to the state of the art without relying on structural FLs.

Our contribution has a significant impact on: (1) cost, as we reduce the number of assays needed to extract the same biological information, by removing the dependency on 1 or 2 experimental structural FLs for accurate cell segmentation and (2) possible discoveries in downstream analysis, as we can monitor additional functionally relevant FLs for the exact same cells, thus obtaining a richer description of each cell’s phenotypic response to a perturbation.

## 2. Data

The context of our work is a very flexible live-cell imaging experimental framework, that enables the analysis of a variety of cell lines and FLs reporting on a wide range of cellular processes over time. Furthermore, experimental parameters such as temporal and spatial resolution can be changed, as well as acquisition conditions. Those characteristics impact the microscope’s acquisition mode and thus the resulting noise and appearance of cells.

Accordingly, it is important to ensure our computational workflow is sufficiently flexible to accommodate changes in FLs, cell lines and acquisition parameters.

### 2.1. Image data

We acquired data on five multi-color reporter cell lines derived from commonly used U2OS [11] and A375 [12] cancer cell lines, that we designated CL1, …, CL5. In this paper, we work with live-cell imaging data where fluorescent labeling is done by tagging proteins of interest with fluorophores, the combination of which we refer to as *fluorescent reporter proteins* (FRPs). Each reporter cell line expresses 3 or 4 spectrally distinct FRPs. Unless otherwise stated, we acquired all images with a Nikon A1R confocal microscope as a live video microscopy imaging sequence. For each experimental condition, we acquired images on 3 *xy* positions every 3 hours for 72 hours, at various optical zooms.

Table 1 helps understanding the structural nature of the non-structural FRPs used in the cell lines evaluated here. While some are structurally strong, labeling consistently a specific organelle (e.g. P1 of CL1), most are dynamic and can be considered as weak. Cell line CL5, for example, only expresses structurally weak FRPs.

**Table 1.**
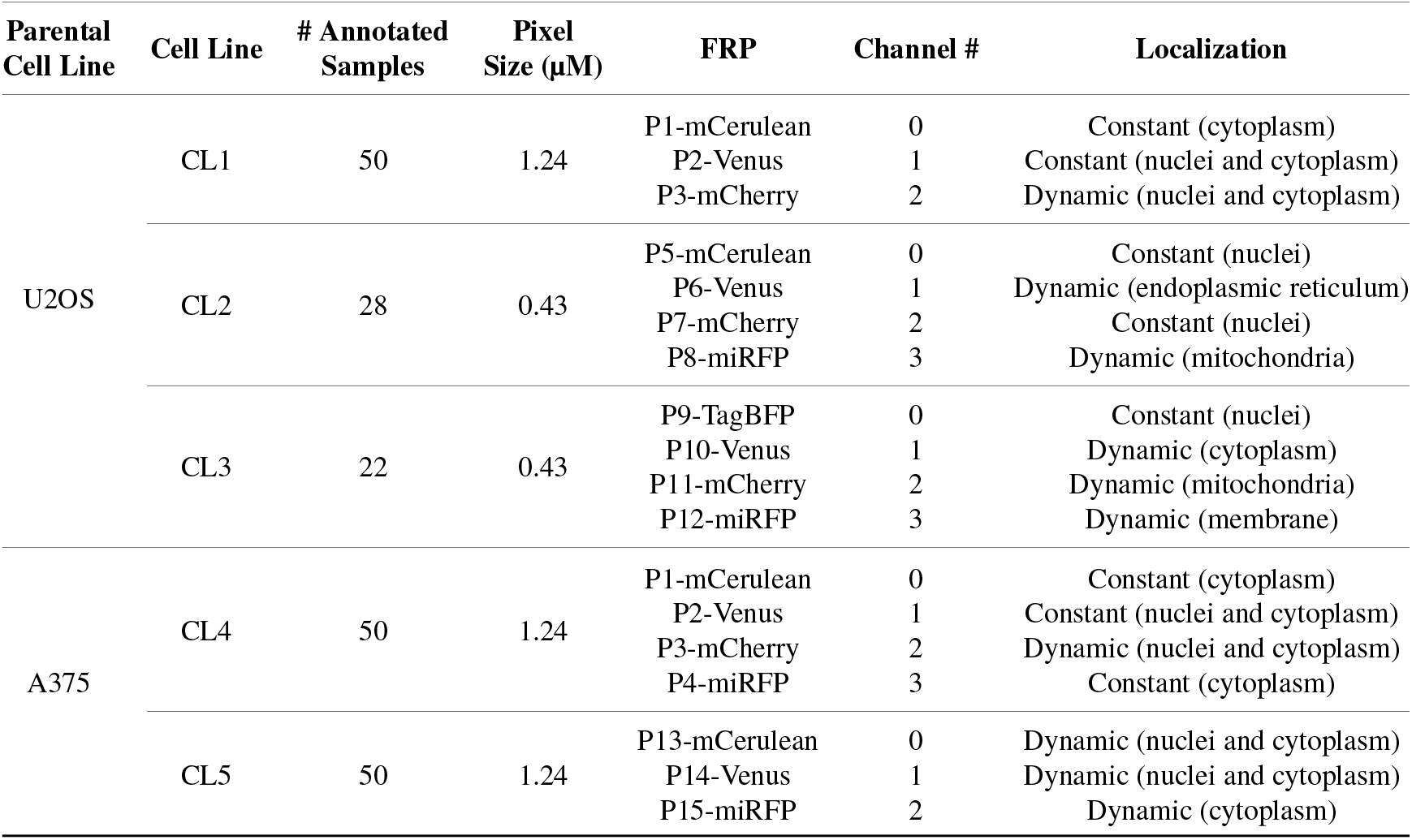
Description of cell line components used in the assays - Parental Cell Line, Cell Line name, Number of Annotated images, Size of pixels, Fluorescent Reporter Protein (FRP), Channel Number and localization. FRPs are tagged to different proteins (P) – in the N or C terminus (FP) – integrated in each of the cell lines.

### 2.2 Annotated dataset

We manually annotated the nuclei and cell boundary of 50 (resp. 28 and 22) images for each cell line/FL combination of cell lines CL1, CL4, CL5 (resp. cell lines CL2 and CL3). We randomly selected these images from different experimental conditions and evenly distributed over time in order to capture the dynamic localization of certain proteins (e.g. due to experimental treatments and levels of expression at different stages of cell cycle) as well as possible variations in population size caused by cell division or cell death. The number of cells per image ranges from 20 cells to more than 100 in some images.

For each set of annotated images, 80% were used for training, 10% for validation and the last 10% for evaluation. With an average of 50 cells per images, we have about 250 cells per validation/test set and approximately 2000 individual cells in the training set, an appropriate number for training and evaluation. Example images and manual annotations are displayed in Figure 1.

**Figure 1.**
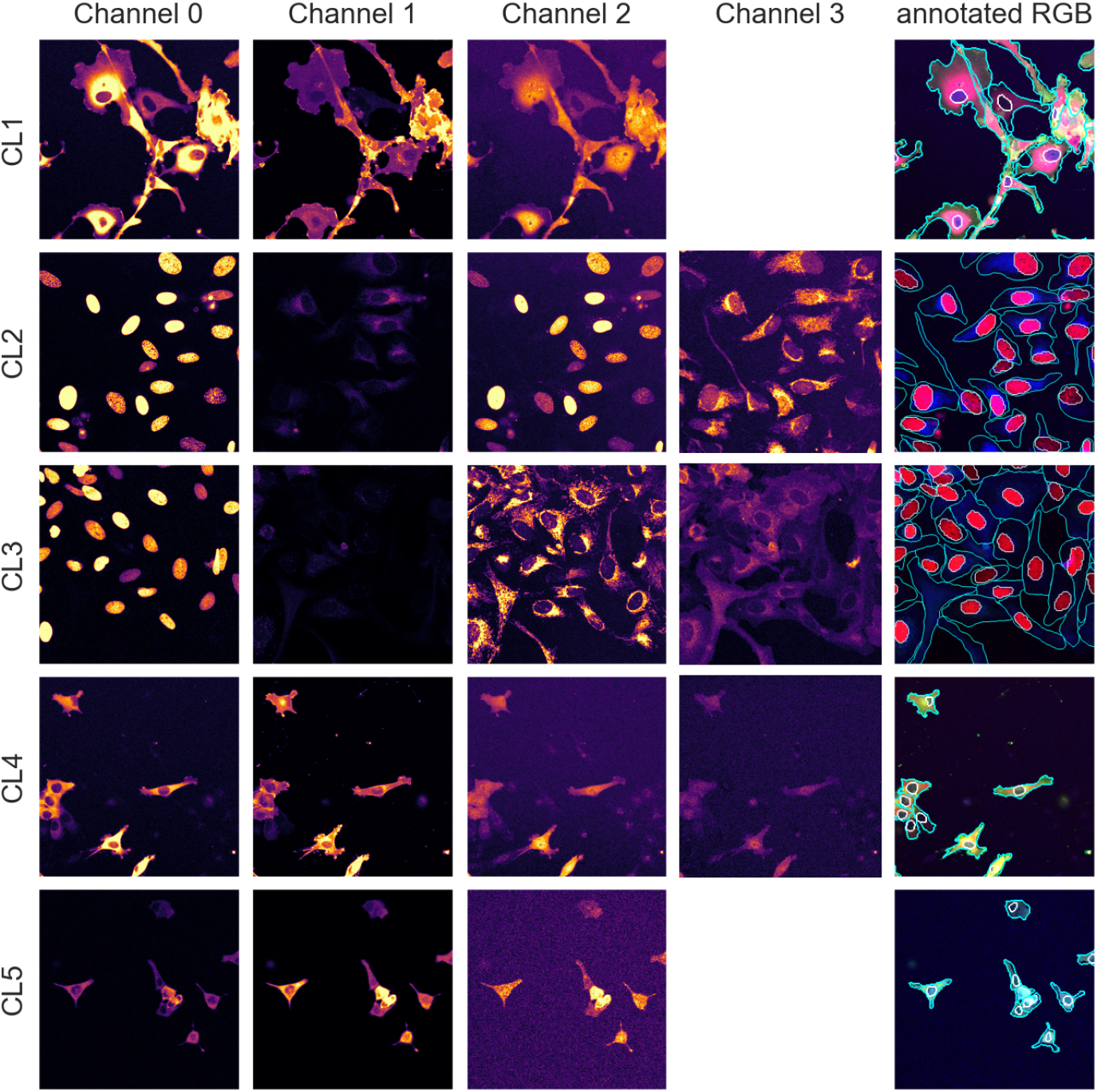
Image samples from the different assays showing individual fluorescence channels as well as a color version with manual segmentation annotations overlay. The images are cropped for ease of visualization.

Furthermore, to test our method for robustness to changes in assay and acquisition parameters, we also annotated 5 additional images of the CL1 cell line acquired with a different microscope (widefield) and higher temporal resolution, resulting in noisier images. We used this dataset for evaluation purposes only as presented in Section 4.3.2.

All datasets are publicly available at doi.org/10.6084/m9.figshare.21702068.

## 3. Methods

### 3.1. Generalist Segmentation Model Backbone

In this paper, we use Cellpose [4], a state-of-the-art cell segmentation model, as a generalist segmentation model backbone. We note however, that our approach is model-agnostic and could therefore use alternative backbones [2], [3].

Cellpose segments images using a three-part pipeline: Firstly, it resizes the images so that the average cell diameter of the dataset conform to the model original training cells’ diameter. Secondly, for a cell object *o* ∈ {*nuclei, cyto*}, a Cellpose model *M*_*o*_ maps a rescaled image *Î* intrinsic intensity space to a flow and probability space (*F*_*X*_, *F*_*Y*_, *P*). The flow maps *F*_*X*_, *F*_*Y*_ are the derivatives (along the X and Y axes) of a spatial diffusion representation of individual cell pixels from the cell’s center of mass to its extremities. Thirdly, Cellpose combines the (*F*_*X*_, *F*_*Y*_, *P*) flow and probability maps to predict instance segmentations *S*_*o*_ using flow analysis and thresholding on all three maps combined. First, the flows *F*_*X*_ and *F*_*Y*_ are interpolated and consolidated where the pixel-wise probabilities *P* are above a pre-set threshold. The instance masks are then generated by analysing the flows histogram from their peak. Cellpose overall segmentation process is summarized in the following equation:

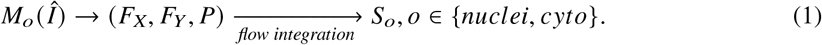

### 3.2. Segmentation Model Finetuning Approach

In order to segment images with non-structural FLs, we leverage the generalization powers of Cellpose to build a model zoo of pre-trained models finetuned on each of our cell line’s FLs using annotated data. Similarly to the out-of-the-box pre-trained generalist Cellpose model (which we call *Vanilla Cellpose* in the remainder of this article), our method rescales the images to the average diameter of the training set.

To finetune Vanilla Cellpose, we train for each organelle and cell line (*o, cl*) combination (with *o* ∈ {*nuclei, cyto*} and *cl* one of the dataset cell lines) several Cellpose models on subsets of channels 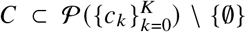 taken from the powerset of the set of channels (excluding the empty set), with *K* the total number of channels. For simplicity we denote this powerset as {*c*_0_, …, *c*_*K*_ }. When |*C* | > 1, each individual channel of *C* from the same image sample are inputted as independent training samples. Each finetuned model *M*_*o,cl,C*_ is then used to predict segmentation flows by evaluating each channel from *C* individually. Those segmentations are then fused together at inference to produce a single segmentation map. To segment an image for an (*o, cl*) combination we end up using the model trained using the channels 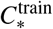 for which the highest score is achieved on our evaluation set using the channels 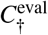.

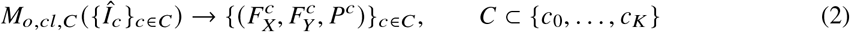

When |*C* | = 1, we refer to the finetuning method as *channel-wise* (CW), such that a model is trained for each individual channel (see upper part of Figure 2). The respective CW models can be used on their respective training channels or can be aggregated together to produce a channel-wise segmentation for several channels at once.

**Figure 2.**
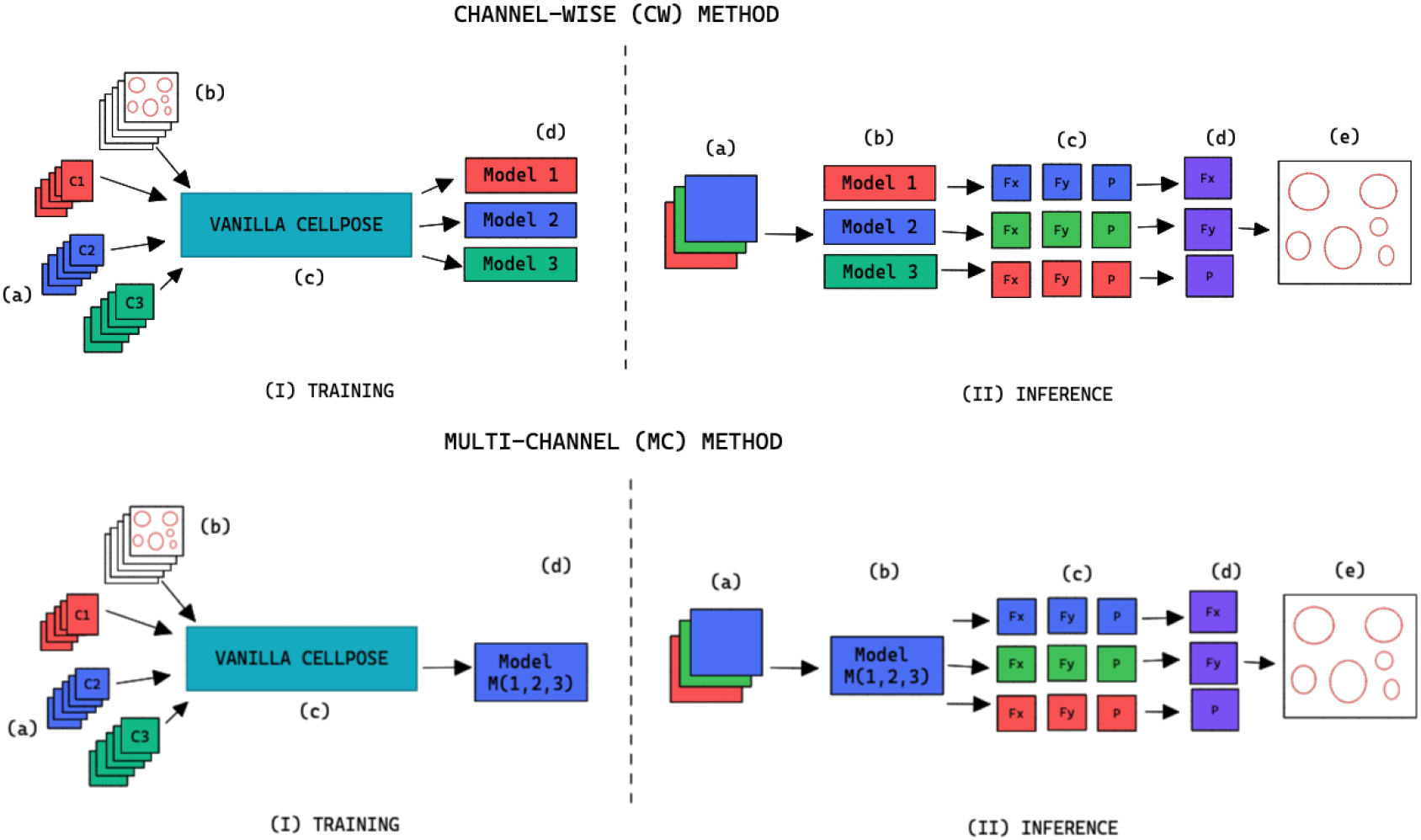
Training and inference workflow for the segmentation of cell organelles without the use of structural FL using the channel-wise approach (top) and multi-channel approach (bottom). **(I) Training:** (a) Training set of multi-modal fluorescent images (3 channels represented as red blue and green), (b) Training set annotations of the organelles segmentations, (c) Out-of-the-box pre-trained Cellpose model (Vanilla Cellpose), (d) Finetuned model trained for each of the individual channels (channel-wise) or trained with a subset of the channels (multi-channel). **(II) Inference:** (a) Multi-modal fluorescent image (3 channels), (b) Models selected from the model zoo corresponding to the image’s cell line and FL channel combination, (c) Spatial flows and probability maps output by the finetuned models for each of the channels, (d) Channel-wise averaging of the maps, (e) Integration into the segmentation labels.

When |*C* | > 1, we refer to the finetuning method as *multi-channel* (MC), such that a model is trained on several channels at once (see bottom part of Figure 2). The MC segmentation models can be evaluated on any subset of the channel set *C* they were trained with.

### 3.3. Segmentation Model Finetuning Parameters

To finetune Cellpose, we train from the available generalist pre-trained model on our dataset using the approach detailed in Section 3.2. We retain the model’s hyper-parameters from the original Vanilla Cellpose training with the following two exceptions:(1) as detailed in Section 3.4, we use non-deterministic augmentations on each of our training samples; (2) we stop the training using early stopping on the validation set [13], with a patience of 50. Both additions are efficient regularization methods limiting over-fitting and contributing to the overall robustness of the segmentation methods with respect to changes in the imaging setting.

### 3.4. Data Augmentation

Augmentation methods are used during training the Vanilla Cellpose model to both virtually increase the size of our dataset as well as offer better generalization. They are performed iteratively from scratch on each image batch. For methods involving random distributions, the parameters are uniformly sampled from a pre-defined parameter range. Each augmentation has an application probability *p*_augment_ = 0.5, adding more variability across epochs and samples. The augmentation methods are described in Table 2.

**Table 2.**
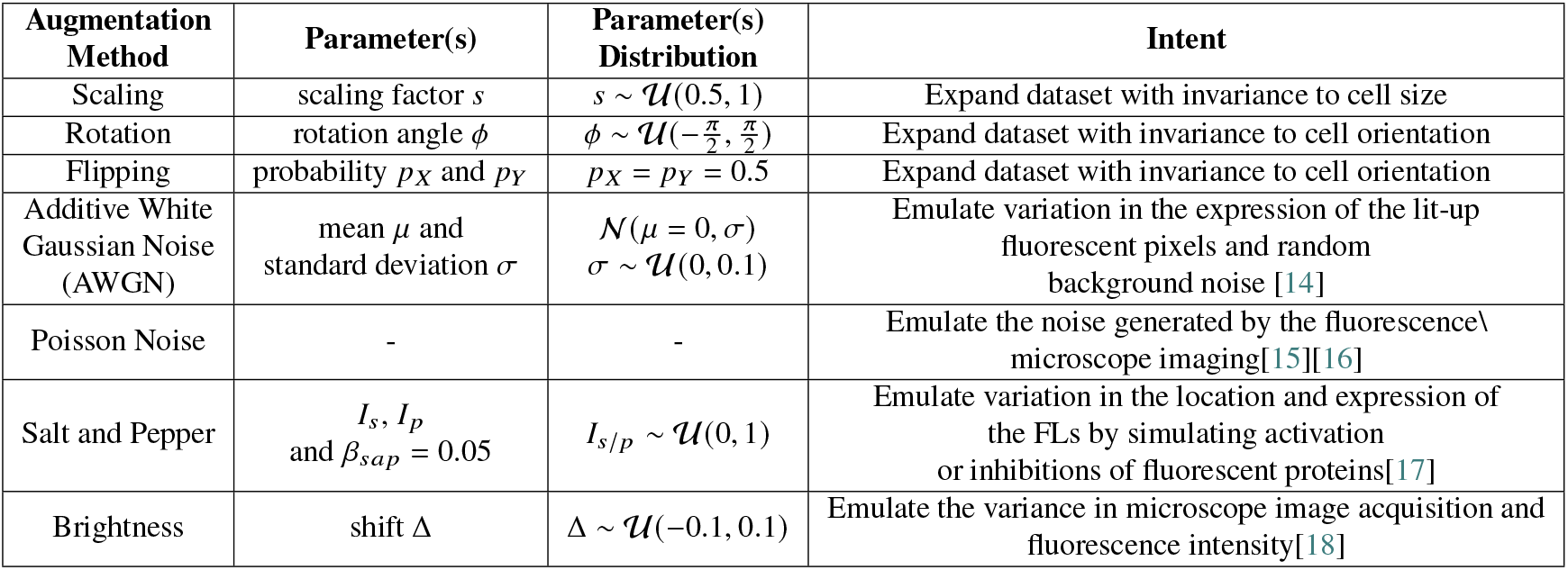
Augmentations applied to the training set during the finetuning of Cellpose models.

| Augmentation Method | Parameter(s) | Parameter(s) Distribution | Intent |
| --- | --- | --- | --- |
| Scaling | scaling factor $s$ | $s \sim \mathcal{U}(0.5, 1)$ | Expand dataset with invariance to cell size |
| Rotation | rotation angle $\phi$ | $\phi \sim \mathcal{U}(-\frac{\pi}{2}, \frac{\pi}{2})$ | Expand dataset with invariance to cell orientation |
| Flipping | probability $p_X$ and $p_Y$ | $p_X = p_Y = 0.5$ | Expand dataset with invariance to cell orientation |
| Additive White Gaussian Noise (AWGN) | mean $\mu$ and standard deviation $\sigma$ | $\mathcal{N}(\mu = 0, \sigma)$<br>$\sigma \sim \mathcal{U}(0, 0.1)$ | Emulate variation in the expression of the lit-up fluorescent pixels and random background noise [14] |
| Poisson Noise | - | - | Emulate the noise generated by the fluorescence\ microscope imaging[15][16] |
| Salt and Pepper | $I_s, I_p$<br>and $\beta_{sap} = 0.05$ | $I_{s/p} \sim \mathcal{U}(0, 1)$ | Emulate variation in the location and expression of the FLs by simulating activation or inhibitions of fluorescent proteins[17] |
| Brightness | shift $\Delta$ | $\Delta \sim \mathcal{U}(-0.1, 0.1)$ | Emulate the variance in microscope image acquisition and fluorescence intensity[18] |

### 3.5. Segmentation Fusion

Once the segmentations are generated for each image channel using finetuned models, a final image segmentation is generated by fusing the individual channel segmentation maps. We implement a method we name *Flow Averaging* (FA) to do so. We also consider several state-of-the-art methods. Fusion methods are described in Table 3.

**Table 3.** Description of the segmentation fusion methods considered to generate aggregated segmentations from channel-wise segmentations

| Method | Description | Reference |
| --- | --- | --- |
| Flow Averaging (FA) (This study) | Fusion of flow maps followed by flow analysis | - |
| Selective and Iterative Method for Performance Level Estimation (SIMPLE) | Iterative majority voting over propagated segmentations, weighted by estimated performance | [19] |
| Simultaneous Truth and Performance Level Estimation (STAPLE) | Statistical fusion framework using hierarchical models of rater performance | [20] |
| Voting (V) | Pixel-wise voting | [21] |
| Majority Voting (MV) | Majority label voting in images patches | [22] |

The FA method uses Cellpose internal representations to aggregate the segmentation maps. FA averages the segmentation probability maps and flow maps obtained by running finetuned models on each channel individually, to obtain a final aggregated segmentation map.

Given |*C* | channels selected for an image, our approach yields |*C* | segmentation tuples 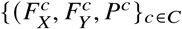 generated by the Cellpose-U-Net(s). From these tuples we average each individual maps along the channel dimensions, yielding maps 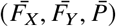. The averaged maps are then transformed into instance segmentation masks using Cellpose’s integration method.

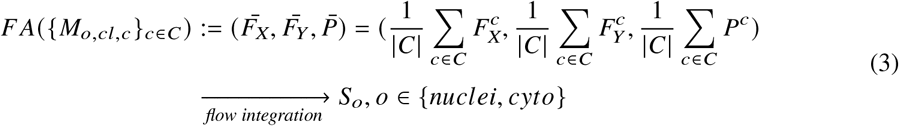

### 3.6. General Workflow Summary

Our workflow can be summarized as follows, for an organelle and cell line (*o, cl*) combination:

1. Annotate the organelles *o* on *N* images of the imaging assay of cell line *cl*
2. Finetune a Vanilla Cellpose model 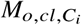 using the FL channels of *C*_*i*_, for all (non-empty) subsets *C*_*i*_ of the powerset of available channels in *cl*. For better readability, we consider the cell line *cl* and organelle *o* fixed in the latter and denote the model 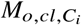 as 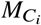.
3. Evaluate the finetuned models on a validation set, using the channel-wise and multi-channel strategies on individual channels or the fusions of channels.
4. Select out the best performing model *M*_*C*\*_ and channels upon which to evaluate it *C*^†^ using an evaluation metric *S* (as defined in Section 4.1) on a validation set:

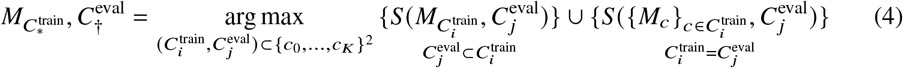

with 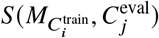 corresponding to the score obtained on average on a validation set when using a model trained on training channel(s) 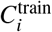 for inference on evaluation channel(s) 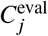, using segmentation fusion when 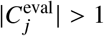.
5. Use 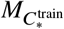 for inference with channels 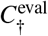 – using segmentation fusion if 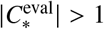 – on new images of the (o, cl) combination, while keeping other models in the model zoo for potential future cell lines stemming from the same parental cell-line and with some intersections in the respective sets of FLs.

## 4. Performance Evaluation Results

### 4.1. Evaluation Metrics

For segmentation evaluation, we use the precision, recall and F1-score. A predicted segmentation is considered as a true positive if the intersection-over-union (IoU) between this segmentation and a ground truth segmentation is above a threshold here set at 0.5. Recall is the percentage of cells detected, precision the probability that a detected cell is really a cell and F1-score is the harmonic mean of the two. We use the F1-score as our principal metric of evaluation for our models performance, as it measures accuracy through both precision and recall. Additionally, we also compute the Jaccard similarity [23], the aggregated Jaccard index [24] and the average precision [2]. The evaluation using these metrics are available in the supplementary material.

### 4.2. Segmentation Fusion Methods Evaluation

Table 4 conveys the performance of the different segmentation fusion methods tested in this work. The benchmarking is shown on the aggregation of all channels for each image, on both nuclei and cytoplasm, using the channel-wise method. The results clearly indicate a better performance when using the Flow Averaging (FA) method introduced in Section 3.5: FA always appears as the best fusion method among the ones benchmarked here, in some cases with a large margin. We believe that this result is owed to the fact that the diffusion maps are optimized representations combining information about the pixel-level probability and object-level shape properties. This makes them particularly useful for fusion and conveys them an advantage over raw image fusion and fusion of segmentation masks. We therefore set FA as the default fusion method, and present our worklow results using FA in the remainder of the paper.

**Table 4.** Comparison of the performance of the different channel fusion methods on the test set images, assessed with F1-score as a segmentation metric. It must be noted that the comparison is drawn between the fusion of all channels for each cell lines, evaluated using the Channel-wise strategy - and not the best scoring strategy or set of training/evaluation channels. Although the fusion method performance holds for other variants of our overall method.

| Organelle | Fusion Method | CL1 | CL2 | CL3 | CL4 | CL5 |
| --- | --- | --- | --- | --- | --- | --- |
|  |  | 0, 1, 2 (CW) | 0, 1, 2, 3 (CW) | 0, 1, 2, 3 (CW) | 0, 1, 2, 3 (CW) | 0, 1, 2 (CW) |
| Cytoplasm | FA | 0.8135 | 0.5138 | 0.3264 | 0.7769 | 0.6873 |
|  | SIMPLE | 0.729 | 0.4744 | 0.2967 | 0.6846 | 0.4218 |
|  | STAPLE | 0.8043 | 0.4043 | 0.2876 | 0.6881 | 0.6635 |
|  | Voting | 0.729 | 0.4571 | 0.3094 | 0.6916 | 0.4218 |
|  | Majority Voting | 0.729 | 0.4344 | 0.3122 | 0.7035 | 0.4218 |
| Nuclei | FA | 0.904 | 0.6871 | 0.843 | 0.8758 | 0.7994 |
|  | SIMPLE | 0.8016 | 0.3804 | 0.7634 | 0.7303 | 0.638 |
|  | STAPLE | 0.8016 | 0.4398 | 0.7341 | 0.6822 | 0.6449 |
|  | Voting | 0.8016 | 0.2968 | 0.7194 | 0.6784 | 0.6449 |
|  | Majority Voting | 0.8016 | 0.3266 | 0.6941 | 0.6817 | 0.6449 |

### 4.3. Segmentation Evaluation

#### 4.3.1. Results

Figures 3 and 4 display the performance of our method using the F1-score metric. Those results were computed using 5-fold cross validation on annotated datasets. We evaluate the performance of Vanilla Cellpose against our method across every combination of training channels (channel-wise and multi-channel) upon every combination of evaluation channel aggregated together using the FA fusion method. The Vanilla Cellpose scores are evaluated on several channels at once using this fusion method as well. Examples of our segmentation are displayed in Figure 5.

**Figure 3.**
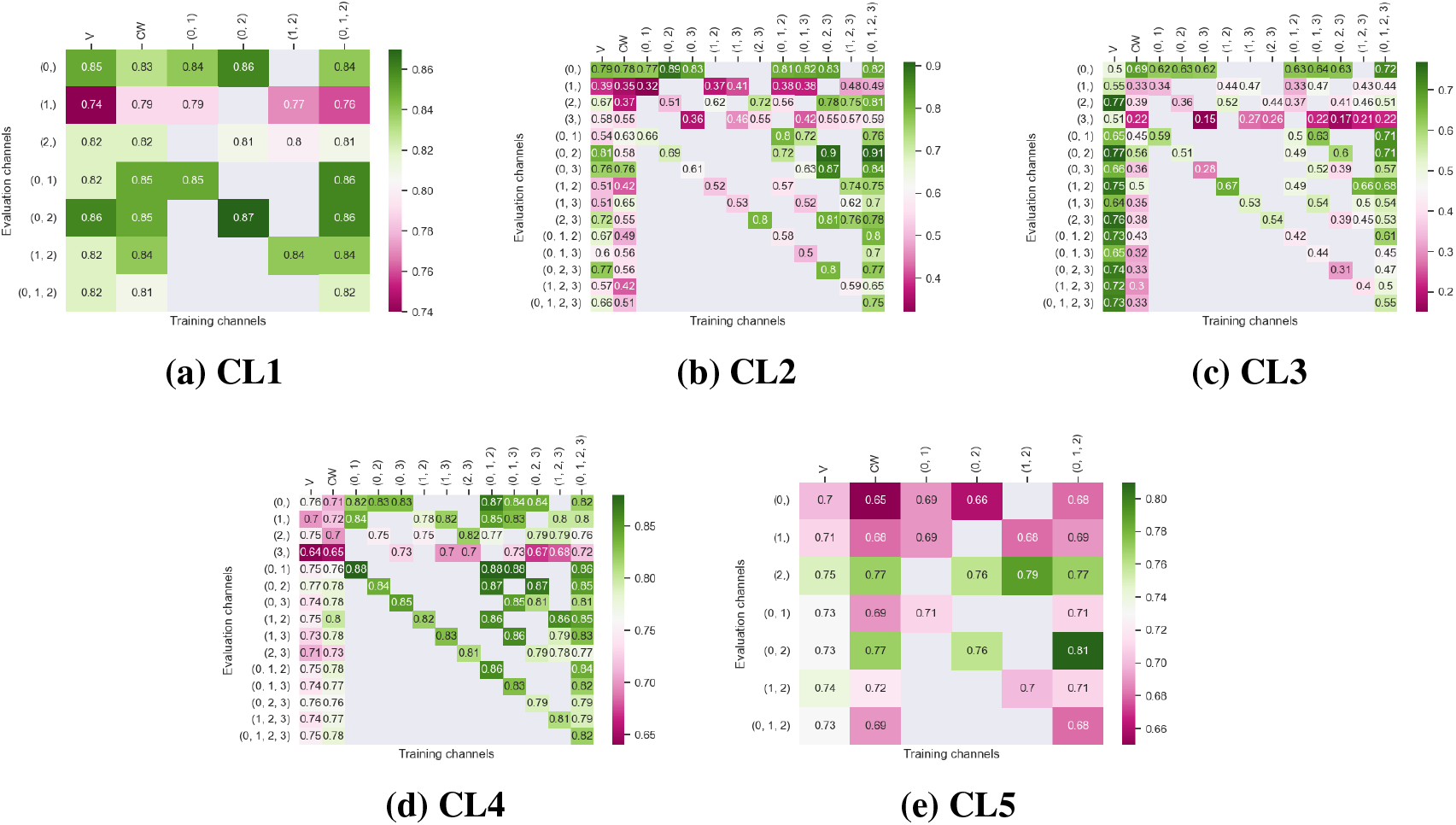
5-fold cross validated F1-scores for cytoplasm segmentation on all 5 cell-lines. These tables show the evaluation using Vanilla Cellpose (V), the channel-wise (CW) strategy and the multi-channel (MC) strategy as columns, on the powerset of channels as rows, aggregated together using the flow averaging (FA) method.

**Figure 4.**
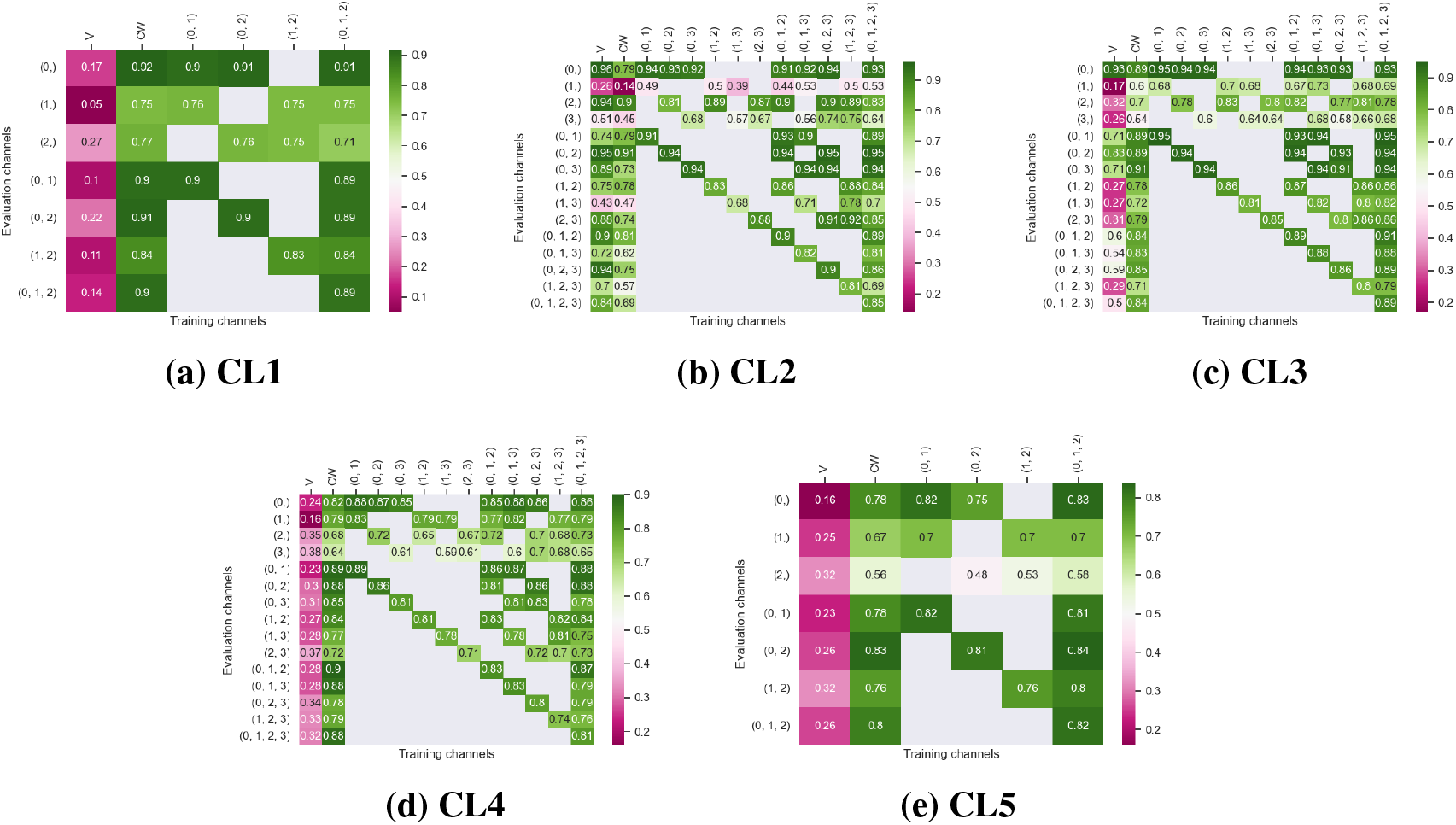
5-fold cross validated F1-scores for nuclei segmentation on all 5 cell lines. These tables show the evaluation using Vanilla Cellpose (V), the channel-wise (CW) strategy and the multi-channel (MC) strategy as columns, on the powerset of channels as rows, aggregated together using the flow averaging (FA) method.

**Figure 5.**
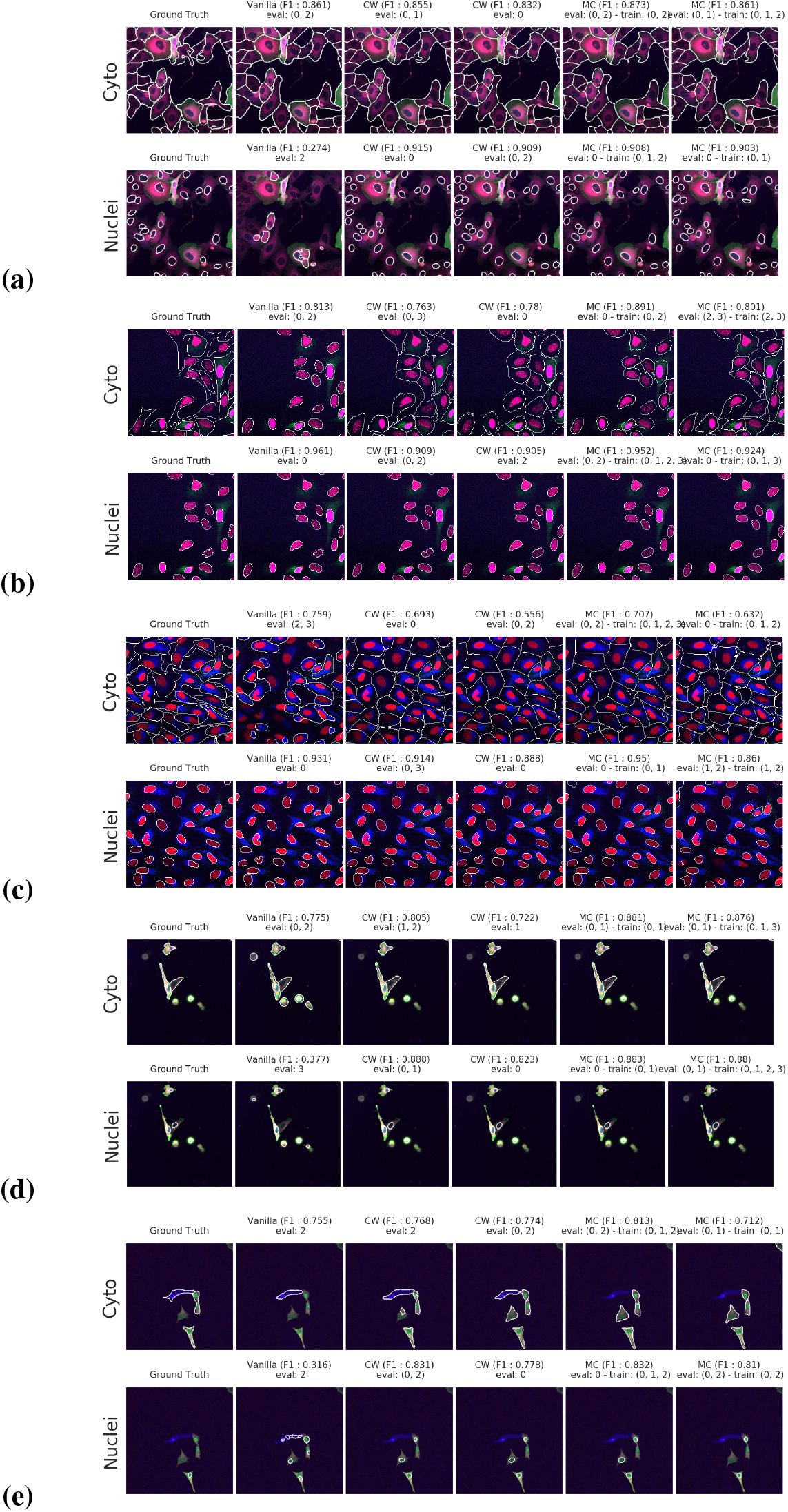
Segmentation examples using the proposed method on the test set images for (a)-(e) CL1 to CL5. We compare: ground truth, Vanilla Cellpose result for best evaluation channel combination, channel-wise (CW) and multi-channel (MC) finetuning strategies. The respective training channel combination and evaluation channel combination are detailed on the figures. The images are cropped for ease of visualization.

Our results indicate that fine-tuning is an essential step when dealing with datasets which do not contain cytoplasmic and nucleic structural FLs, as indicated in Table 1. With fine-tuning, we indeed obtain state-of-the-art level results on individual channels trained independently (CW), the fusion of channels evaluated through independently trained models (CW) and on the fusion of channels trained together (MC). Furthermore, we observe that combining the different segmentations together outperforms not only the Vanilla Cellpose results but also the results of those finetuned models on their respective channels in all cases (the only exception being CL3 on cytoplasm which we will discuss in the next Section). That is particularly striking for structurally weak FLs which, when aggregated together using either the CW or MC approach, reach the segmentation quality of structurally strong FLs (e.g. static FLs in a single organelle). Additionally it can be noted that when trained on a set of channels which include both strong and weak FLs, the model performs well upon evaluation on the subset of its training channels which excludes the structurally strong FLs.

#### 4.3.2. Impact of acquisition method on model generalization

We test the generalization capabilities of our model to other imaging acquisition procedures by evaluating the performance drift of a model trained on images with acquisition parameters *A*_1_ (confocal microscope, image size 512×512 pixels, and image scale of 1.24μM per pixel) when applied to images acquired with parameters *A*_2_ (widefield microscope, image size 2044×2044 pixels, and image scale of 0.32μm per pixel). We use 50 annotated images acquired under conditions *A*_1_ of cell line CL1 for training and testing a model, and 5 annotated images acquired under condition *A*_2_ of the same cell line to evaluate the aforementioned model performance drift. The segmentation evaluation results are shown in Figure 6 and demonstrate the ability of our model to generalize to acquisition method *A*_2_ by producing similar segmentation scores and out-performing Vanilla Cellpose. These results indicate that our approach does not merely finetune Cellpose to our dataset, but rather to a specific set of FLs of a cell line, successfully generalizing to other datasets with the same assay-related conditions.

**Figure 6.**
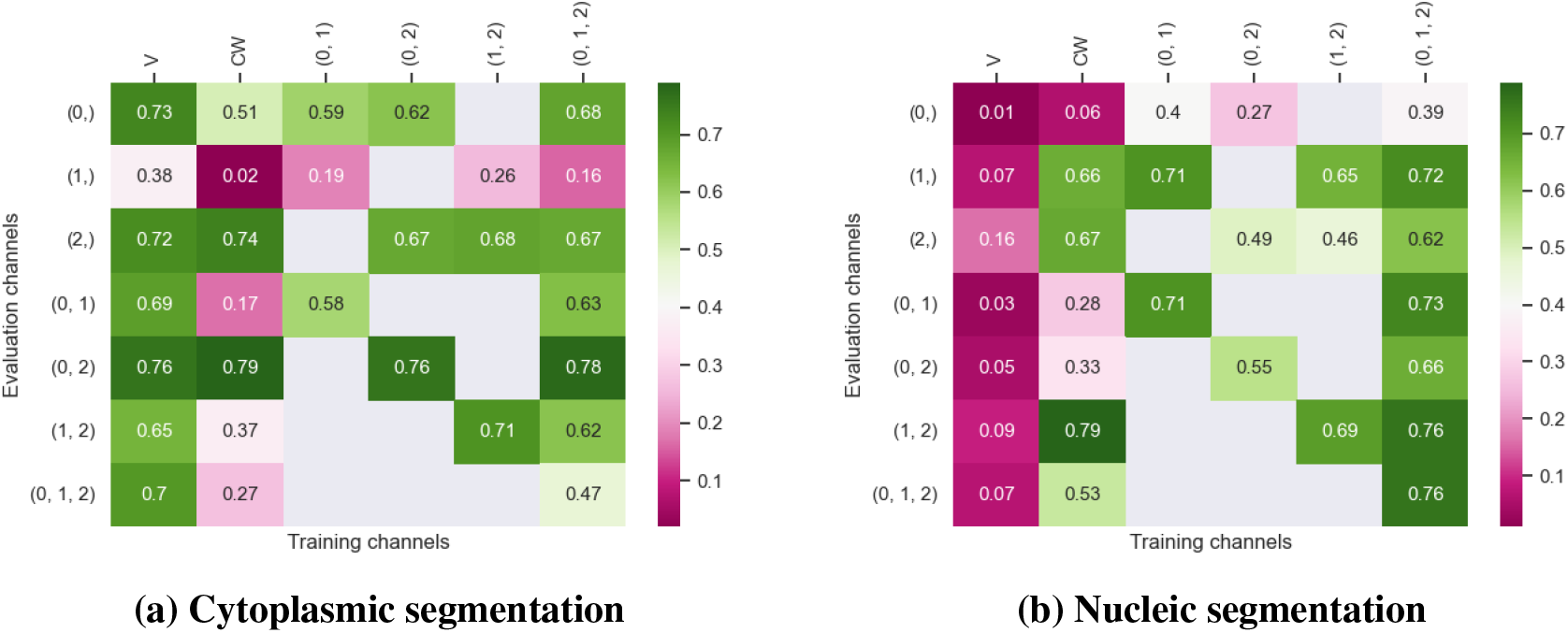
Evaluation of the F1 scores on the CL1 cell-line imaged with the A_2_ acquition method (widefield). The tables are organized the same way as in Figures 3 and 4.

## 5. Discussion

Our work provides a reliable, re-usable and scalable method for the segmentation of cell images without structural FLs, with manageable annotation effort. Our results show that the proposed workflow leads to models outperforming Vanilla Cellpose on datasets with only non-structural FLs, while requiring few annotated examples by leveraging Cellpose extended pre-training.

Our results show that leveraging non-structural (and even structurally weak) FLs in concert improves segmentation, even when the signal is very heterogeneous between cells, and some cells do not appear at all in some channels. Indeed, each channel provides some useful and potentially complementary information on nucleus and cytoplasm, which can be combined by segmentation fusion. Thus, aggregating channels together allows benefiting from complementary non-structural FLs to outline individual cell objects. This is demonstrated by results in Figures 3 and 4 and qualitative examples in Figure 5. This observation is especially salient for cell lines which do not contain any structurally strong FLs (such as CL2 and CL3 for cytoplasm, and CL4 for nuclei) or for cell lines which segmentation models were trained and evaluated without their structurally strong FLs (e.g. CL1 without channel 0 for both nuclei and cytoplasm, or CL2 and CL3 without channel 0 and 2 for nuclei). For example, while cell line CL5 only expresses structurally unreliable FLs, it can be seen in Figure 3 and 4 that it is able to leverage its different FLs together to produce cytoplasmic segmentation with an F1-score of 0.81 when evaluated on the fusion of channels 0 and 2 using a model finetuned on all of its channels together. Excluding a structurally strong channel from the evaluation results in the same conclusion. For example, nucleic segmentation on CL2 scores an *F*1 of 0.7 when trained using channels 1 and 3 in MC, with FLs that highlight the endoplasmic reticulum and the mitochondria. Using Vanilla Cellpose on the same evaluation channels yields an F1 score of 0.4. Similar results can be observed for CL3 on the same channels.

Furthermore, it is significant to note that the use of a structurally strong FL influences the segmentation quality, even when that FL channel is not used at inference. For example, cell line CL3, which has two structurally strong FLs highlighting the nucleic structure (channels 0 and 2), performs nonetheless very well on segmenting nuclei using only the fusion of channels 1 and 3 when trained using all 4 channels (*F*1 = 0.82). The same can be observed for CL4 on cytoplasmic segmentation: excluding channels 0 and 3 during the evaluation yields an F1-score of 0.85 with a model finetuned on all 4 available channels. This is particularly interesting for the segmentation of cell lines containing subsets of the FLs trained for in our model zoo. Future cell lines could benefit from the multi-channel models trained on some of their FLs as well as stronger – although possibly absent – FLs which would improve the segmentation quality.

However one must select non-structural FLs carefully when applying the proposed workflow. Indeed, some FLs by nature or under the influence of a compound introduced in the assay regimen may be too unreliable for the segmentation task. This is exemplified in our results with channel 1 of CL2 on cytoplasmic segmentation. That FL which is dynamic in the endoplasmic reticulum carries almost no information relevant to the cytoplasmic segmentation by itself, as can be seen in Figure 1. Although models finetuned using this channel benefit from its presence (reaching a 0.91 F1-score on evaluation of the fusion of channel 0 and 2 trained using all 4 channels), models evaluated on it or trained with it in over-proportions perform poorly. If a cell-line was constituted only of FLs of similarly unreliable structural information, our worklow would not be able to segment cells. It cannot – like any segmentation method – segment any organelles out of thin air, but it can leverage structurally unreliable FLs together - with structurally medium or strong FLs when they are available - to reach the quality of segmentation one would get by including functionally structural FLs in the cell lines.

For instance, CL3 is only constituted of structurally weak FLs with regards to cytoplasmic segmentation. The presence of those FLs explains the low F1-score displayed in 3(c). However, as can be seen in Figure 5(c), it nevertheless outperforms Vanilla Cellpose in terms of recall and detection of each individual cell. In this specific instance – with highly dynamic and unreliable FLs – our method generates segmentation masks in the likes of a Voronoi diagram. While not optimal in its boundary detections, it translates to a better segmentation than Vanilla Cellpose, especially in the context of single-cell phenotypic analysis.

Finally, our approach is limited by the same limitations as Cellpose. Although it works as an “expertization” of generalist models similarly to [10], it is still constrained by the same limitations. For example, it does not handle occluded or overlapping cells. It is also susceptible to merging or splitting of individual cell instances which could only be corrected with a robust post-processing step. It is also limited in terms of building cell objects, as – like Cellpose – it detects nuclei and cytoplasm independently, thus yielding standalone cytoplasm and nuclei. In theory, these limitations can be overcome by using a different backbone than Cellpose, one which would handle such issues. It is indeed possible to apply our methodology with different architectures using our finetuning approach, although changes to the fusion method would be required.

## 6. Conclusion

In this work, we have designed, built and tested a workflow to train and infer segmentations of nuclei and cytoplasm on images without functionally structural FLs and for a variety of cell line/FLs configurations. We demonstrated that our approach could be used for a range of assays while freeing up fluorescence channels for two experiment-specific FLs, where the use of structural FLs take space. We have shown that satisfactory segmentation performance can be achieved and replicated on various assays by leveraging multiple non-structural FLs. The advantage of our method is that the freed fluorescence channels can then be used to monitor additional functionally relevant FLs. We thus obtain a richer description of each cell’s response to the perturbation, while limiting costs of assays needed to obtain the same biological information.

Our method is easily adaptable to fit a generalist image processing pipeline and be applied on various assays by aggregating segmentation models trained on multiple cell lines and FLs into a model zoo. Such a zoo of fine-tuned models will greatly support microscopy based cellular assays, and HCS.

## Acknowledgements

We sincerely thank Amritansh Sharma for his detailed and insightful feedback on the segmentation workflow. In addition, we thank Oliver Lai, Claire Repellin and Amritansh Sharma for their help annotating images. Finally, we also thank Catherine Lacayo, Claire Repellin, Oliver Lai and Laura Savy for developing the cell biology tools and conducting the assays that provide the data used in this work.

## Data Availability Statement

The data used in this work can be found at doi.org/10.6084/m9.figshare.21702068.

## Ethical Standards

The research meets all ethical guidelines, including adherence to the legal requirements of the the countries in which the study was conducted.

## Author Contributions

Conceptualization: D.Z.; A.F.; T.W. Methodology: D.Z.; A.F. Formal Analysis: D.Z.; A.F.; T.W. Data curation and visualisation: D.Z. Software: D.Z. ; A.F. Funding acquisition: A.F.; S.R.; T.W.; M.L. Supervision: A.F.; T.W. Writing original draft and reviewing finalized version: all authors.

## Supplementary Material

Supplementary material will be submitted upon acceptance of this manuscript and/or at the request of reviewers. Our supplementary material contains additional results tables with alternative metrics.

